# Precise Traits From Sloppy Components: Perception and the Origin of Phenotypic Response

**DOI:** 10.1101/2022.12.27.522001

**Authors:** Steven A. Frank

## Abstract

Organisms perceive their environment and respond. The origin of perception-response traits presents a puzzle. Perception provides no value without response. Response requires perception. Recent advances in machine learning may provide a solution. A randomly connected network creates a reservoir of perceptive information about the recent history of environmental states. In each time step, a relatively small number of inputs drives the dynamics of the relatively large network. Over time, the internal network states retain memory of past inputs. To achieve a functional response to past states or to predict future states, a system must learn only how to match states of the reservoir to the target response. In the same way, a random biochemical or neural network of an organism can provide an initial perceptive basis. With a solution for one side of the two-step perception-response challenge, evolving an adaptive response may not be so difficult. Two broader themes emerge. First, organisms may often achieve precise traits from sloppy components. Second, evolutionary puzzles often follow the same outlines as the challenges of machine learning. In each case, the basic problem is how to learn, either by artificial computational methods or by natural selection.

## 1. Introduction

Response to an environmental signal requires two steps. First, the signal must be perceived. Second, a response must follow. The evolutionary origin of two-part traits presents a puzzle. Perception without response provides no benefit. Response without perception cannot happen.

Preexisting perceptions or responses may be modified. With a partial step on one side, an evolutionary path opens to solve the new challenge. Modification of prior adaptive traits may be a common pathway.

This article poses an alternative solution. In essence, a purely random preexisting biochemical or neural network within the organism can provide the initial perceptive basis for the evolution of precise responsiveness. If so, then we gain understanding of how organisms may acquire truly novel responsiveness.

In addition, we may begin to understand one of the great puzzles in life. How do organisms acquire a wide array of relatively precise traits given that biological components are inherently stochastic and often unreliable? How does precision arise from sloppiness? Consider perception. We require that external signals induce an internal change in state. To analyze how random systems can acquire and store information, the computational literature has recently built on the idea of liquid state machines.

Think of the smooth surface of a liquid in a container. Drop a pebble on the surface. Waves move across the surface. Drop another pebble, and then another. At any point in time, the pattern of surface waves contains a reservoir of information about the temporal history.

Randomly connected networks act similarly. External inputs enter via sensor nodes. Those signals propagate through the network based on the random patterns of internal connectivity and rules for updating. At any point in time, the network contains information about the temporal history of inputs. The network functions as a dimensional expansion reservoir, transforming time into extent.

A random biochemical or neural network may act as a perceptive internal reservoir. The two-step challenge of perception and response reduces to the much easier problem of evolving an internal response to the perceptive reservoir. It may be possible to achieve an adaptively responsive trait arising from sloppy underlying components.

The remainder of this article provides details. The next subsection gives additional background and references to the computational and biological literature. The following analysis develops a model to illustrate how random networks store information about environmental inputs, creating the basis to predict future environmental states and respond accordingly.

A following subsection speculates that critical learning periods allow individuals to adjust their responses to their unique internal wiring and pattern of reservoir information. The Conclusions consider some possible tests of the ideas and some future directions.

### 1.1. Background and literature

Maass et al.[1] introduced the liquid state machine. The concept, outlined in the introduction, describes a general way in which large dynamical systems retain memory of their past inputs. At any point in time, that memory encoded in the current state of the system can be used to compute responses. The responses may achieve particular goals or predict future inputs.

Computationally, liquid state machines have a recurrent architecture. Roughly speaking, recurrence means feedback loops between internal states [2]. For example, a recurrent computational neural network updates internal states sequentially. External inputs modify the first layer of the network. The first layer then modifies the second layer, which may then modify the third layer, and so on. Recurrent connections flow updates backwards, from a later layer to an earlier layer. Recurrence greatly enhances the computational power of neural networks, in part by storing an internal memory of past inputs.

Recurrent neural networks led to many of the great recent advances in artificial intelligence. However, it can be very difficult to tune the particular connections and dynamic update rules in a network to achieve a particular function.

To solve the tuning problem, one may separate the accumulation of environmental information and memory from the computation of a response to that information. In the simplest application, one can use a randomly connected dynamical system as a reservoir of information and memory about inputs. One can then use a relatively simple computational learning or optimization method to match the current internal state of the reservoir to the desired goal. Often, basic regression methods such as ridge regression are sufficient.

This two-step solution has led to many developments in the computational literature, typically under the topics of reservoir computing or echo state networks [3–5]. Reservoir computing has also grown into a common approach in neuroscience modeling [6], with additional applications using biochemical networks as reservoirs [7,8]. In both computational and neuroscience models, reservoir connectivity patterns other than purely random often arise [5,9–11]. For nonrandom reservoirs, the idea is that particular kinds of information may be better retained by particular architectures. Typically the architectures are not optimized for each application. Instead, a few broad architectural varieties are explored in relation to a particular challenge.

A couple of articles note the potential of reservoirs to help in the understanding of various evolutionary problems [12,13]. My own focus is also evolutionary but limited here to two particular questions. First, can random reservoirs be a potential solution to the puzzle of jointly evolving perception and response? Second, can we place the perception-response problem within the broader frame of precise traits from sloppy components?

## 2. Materials & Methods

### 2.1. Perception and response

The joint evolution of perception and response may be easier if an initially random reservoir can solve the perception side of the puzzle. If random reservoirs provide information that can be the basis for perception, then the evolutionary path to a perception-response system may not be so difficult. In essence, a random system provides sufficient perception to get started and so, initially, only the single response trait must improve evolutionarily to make a workable system. The origin of a workable system provides the opportunity for further evolutionary refinement.

In this article, I limit the analysis to illustrating how random reservoirs provide the capacity for perception and the basis for developing a predictive response. The model brings the key ideas into the evolutionary literature within the context of a simple but important evolutionary puzzle.

The model has three parts. First, environmental inputs come from a chaotic dynamical system. A single parameter of the chaotic system describes the difficulty of predicting future input values. Second, the chaotic environmental inputs feed into a random network that acts as the reservoir. Third, an optimized regression model predicts future input values by using the internal reservoir states as predictors. The quality of the predictions is measured by evaluating additional input data and reservoir dynamics not used in the regression fitting procedure.

### 2.2. Chaotic dynamics

I use the classic Lorenz–96 model for chaotic dynamics [14–16], which is

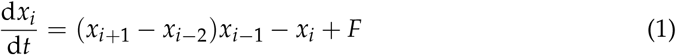

for *i* = 1, …, *N*, with *x*_−*k*_ = *x*_*N*−*k*_ and *x*_*N*+*k*_ = *x*_*k*_, and *F* as the single parameter that describes a constant forcing input. The symmetry of the model means that the long-run trajectories for each dimension have similar properties. I use *N* = 5 for all analyses in this article.

The system tends to be more chaotic as *F* rises above 8 (Fig. 1). Chaos means that a small perturbation at a particular time causes the future system trajectory to diverge from the trajectory of an unperturbed system. The greater the rate of divergence, the less predictable the system.

**Figure 1.**
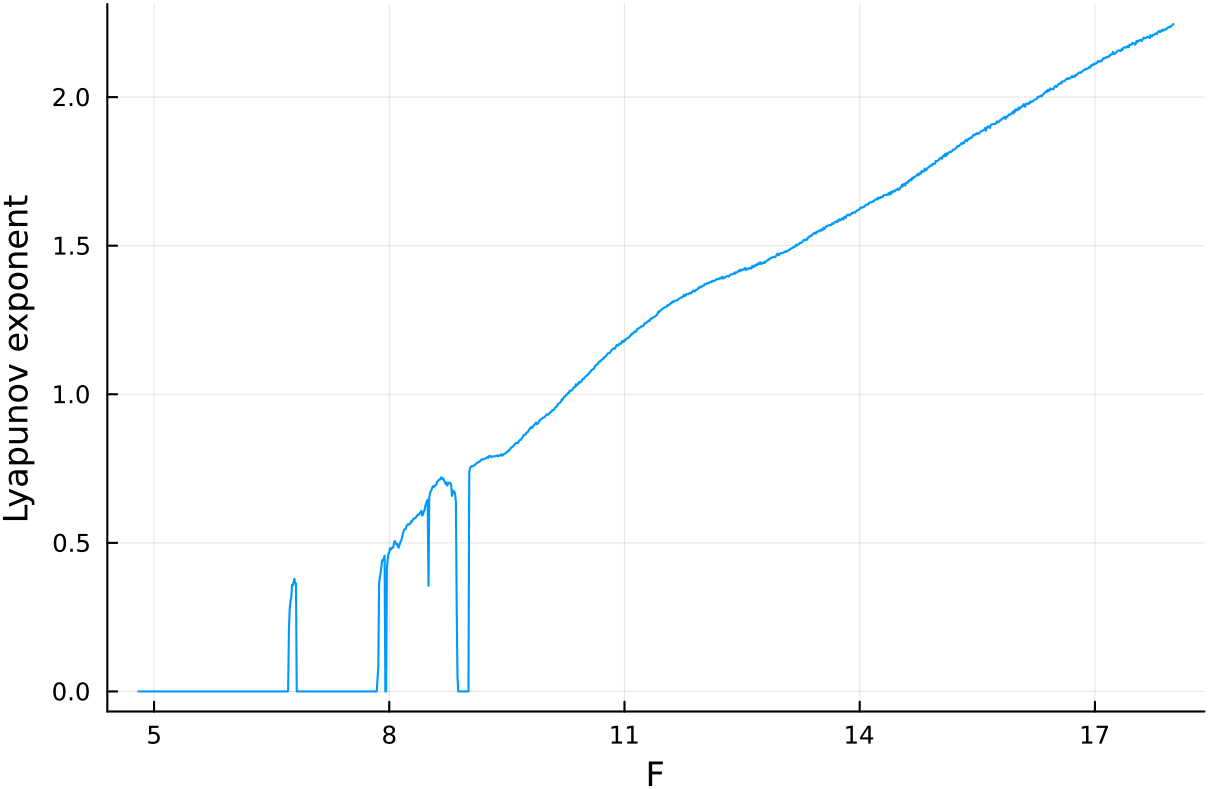
Estimate for the relative speed of chaotic divergence in the dynamics of the Lorenz– 96 equations given in eqn 1, with *N* = 5. Here, the Lyapunov exponent, *λ*, estimates the relative divergence rate. The analysis in this article focuses on the doubling time for divergence, *dbl* = log 2/*λ*, in which a lower doubling time means that future values of the trajectory are harder to predict. For a few limited regions of smaller *F* values, the estimated Lyapunov exponent drops below trend. Those deviations may arise from numerical limitations or a complex pattern of nearly stable periodicity. Sufficiently complex periodicity poses a significant challenge for prediction. The analyses in this article avoid those erratic regions.

Typically, one quantifies the rate of divergence by the dominant Lyapunov exponent, *λ*. Similarly, the system predictability can be quantified by the doubling time of the distance between divergent trajectories, which is *dbl* = log 2/*λ*, with *dbl* denoting a variable. A faster doubling time means that future values of the trajectory are harder to predict. I calculated the dynamics of eqn 1 and the Lyapunov exponent with the Julia package DynamicalSystems [17]. The system becomes increasingly chaotic as *F* rises above 8, which means that *λ* increases and *dbl* (predictability) declines.

### 2.3. Random reservoir

I computed the random reservoir state using the Julia package ReservoirComputing [18]. The reservoir takes the *N* inputs from eqn 1 and updates its *size* internal states. The cited documentation gives the details of the reservoir dynamics architecture and calculations. The outcome arises from the common principles of liquid state machines.

A particular run starts with random initial conditions for the input dynamics and a randomly structured reservoir. Then over the *T* time units of a run, the inputs are feed into the reservoir every 0.01 time units, which triggers an update to the reservoir states. For each of the *T*/0.01 time steps, the reservoir has *size* different state values. Those state values can be used to predict future values of the inputs.

## 3. Results

### 3.1. Predicting future inputs

Briefly, a random reservoir provides sufficient information for the system to predict future inputs of the chaotic environmental dynamics. The more strongly chaotic the system, the shorter the divergence doubling time, *dbl*, and the shorter the time forward for successful predictions. Larger random reservoirs improve the system’s ability to predict future input values. Supporting details follow.

I first calculated the external inputs from eqn 1 at each of the *T*/0.01 time steps, with *T* = 20, 000 for all analyses. I then split the time periods into a training set for the first 0.7*T* = 14, 000 of the time units and a test set for the remaining 0.3*T* = 6, 000 time units. Time units are arbitrary. Predictions provide value if the time extent of predictive success corresponds to a biologically valuable foresight.

Figure 2a shows an example run of the model predictions. The blue curve is the external input value for the first dimension of the Lorenz–96 system, *x*_1_, in eqn 1. The plotted value is rescaled so that the range over the training set is [−1, 1]. The plot shows the final 20 time units of the test set, the time period 19, 980–20, 000.

**Figure 2.**
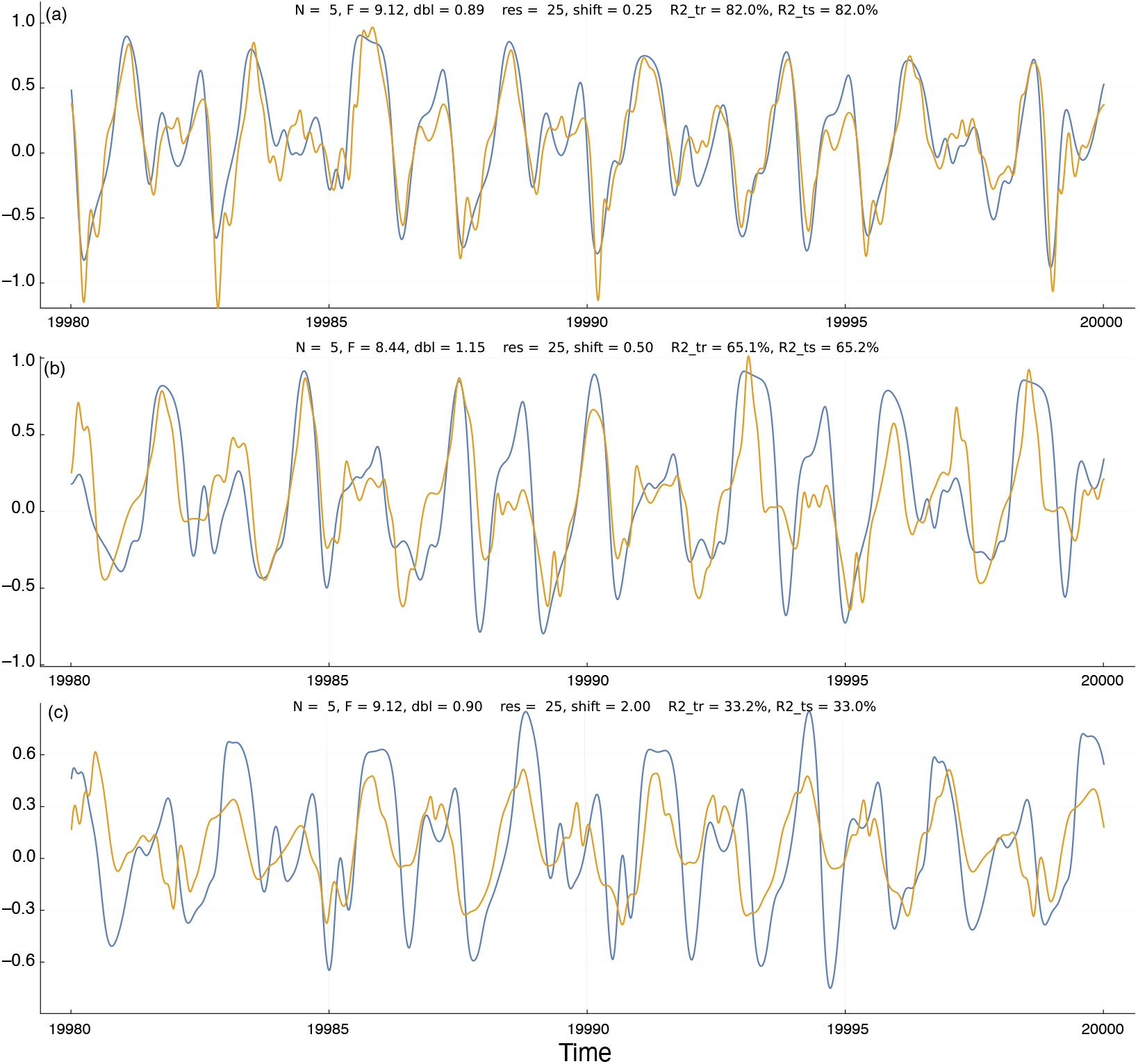
Temporal dynamics of environmental state (blue) and system prediction for the environmental state (gold). At each time point, the internal system uses the information in its reservoir to predict the environmental state *shift* time units into the future. The gold prediction curve is shifted to the right by *shift* time units, so that the closeness of the match between the two curves describes the quality of the predictions. Above each panel, the parameters *N* and *F* describe the environmental dynamics in eqn 1, *dbl* gives the doubling time for the deviation distance of a small perturbation to the dynamics, *res* is the reservoir *size, R2_tr* and *R2_ts* are the R-squared values that describe the percentage of the variation in the blue dynamics curve captured by the gold prediction curve for the training and test periods, respectively, as described in the text. The panels (a), (b), and (c) have corresponding labels on the curves in Fig. 3a. Time units are nondimensional and can be chosen to match the scaling of the environmental process under study. Here, the plots show the 20 time units at the end of the test period of the machine learning procedure used to generate the curves. Here, the abbreviations *res, shift, size, dbl, R2_tr*, and *R2_ts* denote variables. Execution times for the parameters in (b) with reservoir sizes, *res*, of 25, 50, 100 are approximately 58 s, 118 s, 253 s. Timing done on Apple Mac Studio M1 Ultra with Julia 1.9.1, source code git commit a7f74f1. The code was not optimized for execution speed.

The gold curve shows the system’s prediction for future values of the external chaotic input, *x*_1_. For a time point, *t*, the system predicts *x*_1_ at time *t* + *shift*. To compare the predicted input value to the actual input value, I shifted the gold curve by *shift* time units to the right. Thus, each time point on the plot shows the system’s observed and predicted value for time *t*.

I calculated the predicted values by fitting a Bayesian ridge regression model to the training set of observed *x*_1_ values based on the *size* predictors from the internal reservoir states. In Fig. 2, *size* = 25 for all three panels. I obtained the fitted model by the BayesRidge function of the Python scikit-learn 1.2.0 package [19]. I accessed the Python code via the Julia machine learning package MLJ [20].

In Fig. 2, I show the actual input values and predicted input values over the test set of observations. Those test data were not used during the fitting of the ridge regression model and so describe how well the model predictions fit additional observations from the chaotic inputs. I measured the quality of the predictions by the R-squared value, which is the fraction of the variance in the actual input values of the blue curves explained by the predicted input values of the gold curves. For example, the R-squared value for Fig. 2a is 82%, a close fit.

To avoid overfitting the ridge regression model, I used MLJ’s TunedModel function to optimize the BayesRidge hyperparameters for the training period data. That procedure shuffles the data provided for fitting in a way that minimizes overfitting. To test for overfitting on the training data, above each panel in Fig. 2 I show the R-squared values for the training period (*R2_tr*) and the test period (*R2_ts*). The close match of those values demonstrates that the model was not overfit to the training data.

In Fig. 3a the different colored curves show the quality of the predictions for different *shift* time values into the future. The prediction quality on the *y*-axis is given by the R-squared values of the test period. Shorter time shifts into the future provide better predictions, as expected. The *x*-axis shows the doubling time, *dbl*, for trajectory divergence. Greater doubling times correspond to weaker chaotic dynamics and greater predictability. The a,b,c labels on the curves in Fig. 3a match the three panels of Fig. 2. The different panels of Fig. 3 show that increasing reservoir size leads to better predictions.

**Figure 3.**
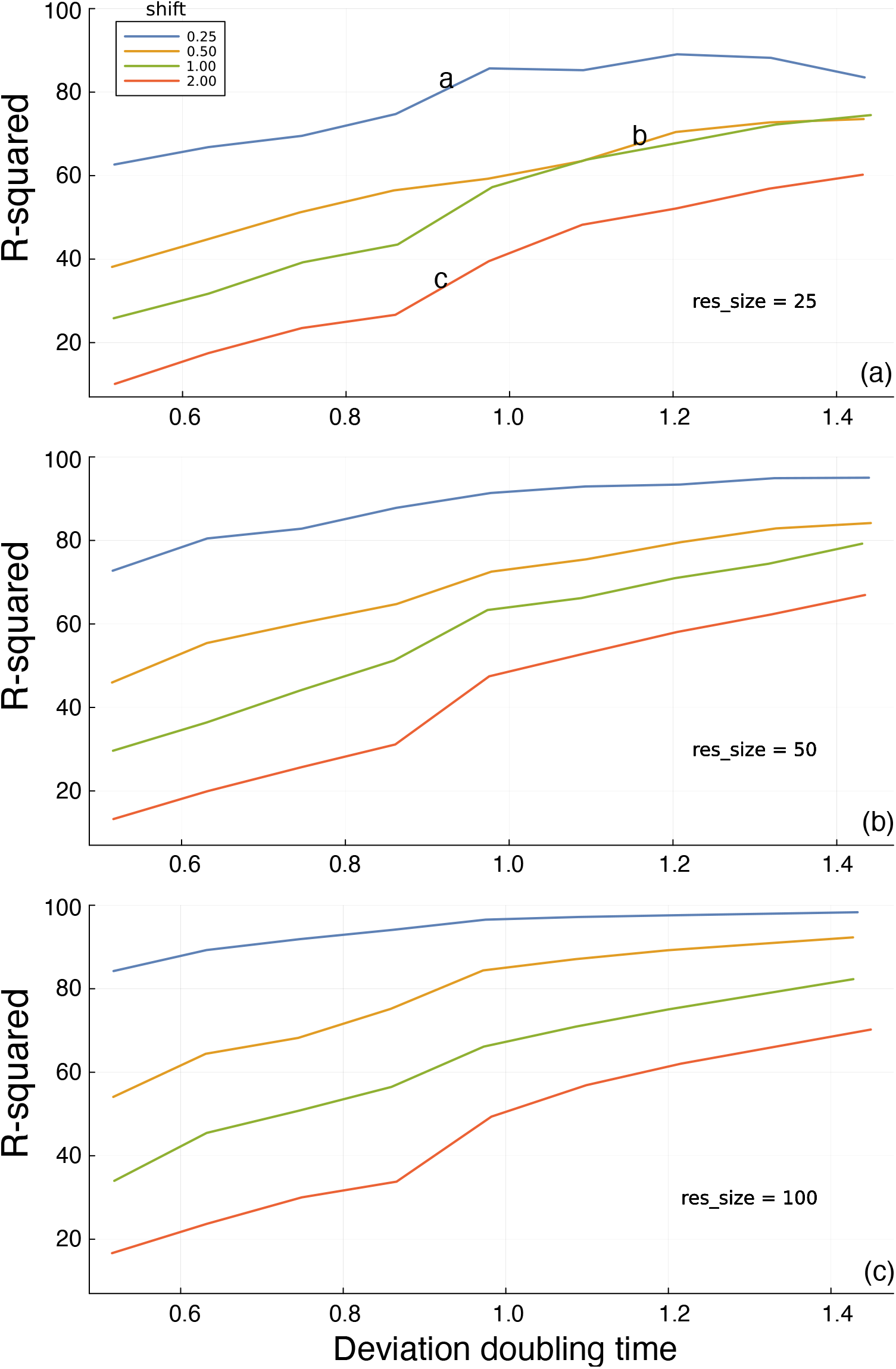
Prediction of future environmental state based on the information in a random reservoir network. Figure 2 shows the environmental dynamics and the prediction challenge. In this figure, the *y*-axis measures the percentage of the total variance (R-squared) in environmental state explained by the predictions generated from the internal reservoir, reflecting the potential for adaptive response. The *x*-axis shows the intrinsic predictability of the environment, measured by the time required to double a small initial perturbation to the dynamic trajectory. The different colored lines describe the time shift into the future at which predictions are compared to actual future dynamics. The *res_size* parameter in each panel gives the *size* of the random reservoir. The a,b,c labels in panel (a) match the corresponding panels in Fig. 2. Each line connects the outcomes at the following 11 approximate doubling times: 0.52, 0.54, 0.58, 0.64, 0.70, 0.77, 0.86, 0.90, 0.99, 1.15, 1.43.

I calculated the test R-squared value *R2_ts* for each parameter combination from one replicate. In Fig. 3, the consistency of the trends across different doubling times and reservoir sizes implies that the variability within a parameter combination is low. If that were not true, then the trends would be much noisier than observed.

To check the actual variability among replicates for a parameter combination, I calculated *R2_ts* for a sample of 20 independent runs for each reservoir size of 25, 50, and 100, using for the other parameters N=5, as in all reported results, F=8.75, corresponding to a doubling time of about 1.0, and a shift value of 1.0.

For any given reservoir size, the variation among samples is small. Reporting results as (minimum, median, maximum) for each set of 20 replicates, the results for reservoir size 25 are (56.0, 57.8, 58.9), for size 50 are (60.0, 61.4, 62.7), and for size 100 are (66.0, 67.1, 68.3). Figure 4 shows that increasing reservoir size improves the prediction of future envi-ronmental state.

**Figure 4.**
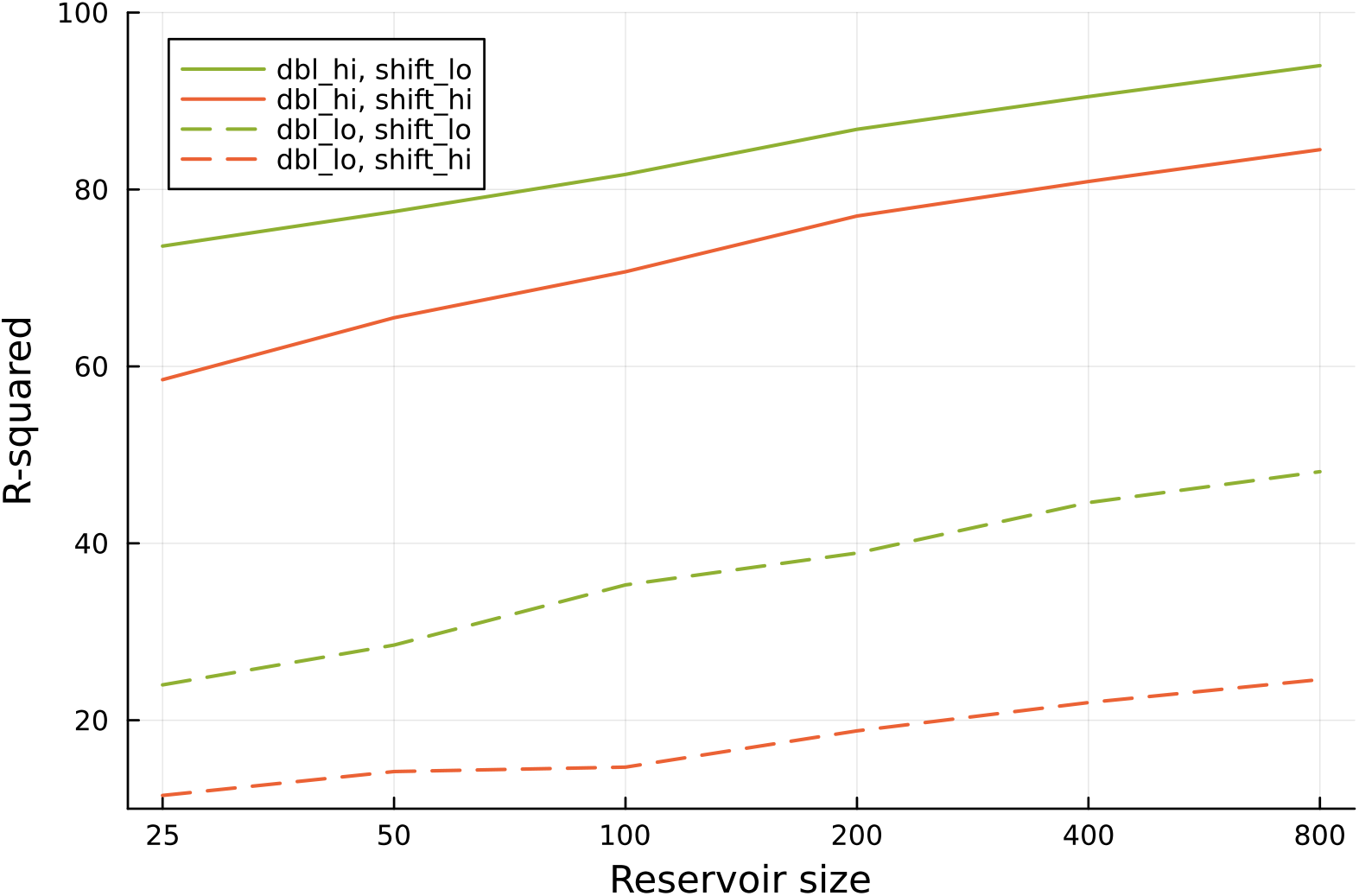
Increasing reservoir size provides better predictions for future environmental state. The analysis follows the methods used in Fig. 3. Here, dbl_lo denotes a doubling time of approximately 0.52, and dbl_hi denotes a doubling time of approximately 1.42. The value of shift_lo denotes a prediction into the future over 1.0 time units, and shift_hi denotes a prediction into the future over 2.0 time units. The six different reservoir sizes used in the computer runs are shown as labels for the tick marks along the *x*-axis.

### 3.2. Critical learning period

The wiring of internal reservoirs may be fixed. For example, the parameters of a simple biochemical network within a cell may be determined primarily by DNA sequence. The network may be random in the sense that it was not shaped by natural selection to capture specific information. But such a random network may be relatively consistent from one individual to another. If so, then the readout of the network to achieve a function may also be fixed among individuals.

Simple neural networks may also be relatively consistent from one individual to another. However, larger networks likely have some stochasticity in wiring. Stochasticity means that random reservoirs of perceptual information may vary from one individual to another. If so, then the way in which individuals read their reservoirs to achieve a function may have to be partially learned.

The demand for such learning may impose the need for a primitive kind of critical learning period in which individuals associate their particular internal reservoir state with successful actions. Such learning periods would be simpler than the kinds of learning that are sometimes observed in the advanced neural systems of vertebrates. Although speculative, the logic for such kinds of critical learning seems compelling.

### 3.3. Other ideas for future study

Comments arising in the review process for this manuscript raised three interesting ideas for future study. First, heritable variation in network size and wiring architecture may provide the opportunity for selection to improve environmental perception. The com-putational literature on reservoir computing provides insight into how different reservoir networks perform with respect to different kinds of environmental challenges [5,18].

Second, environmental change often requires organisms to modify some aspect of their perception or response. In the reservoir model, a change in response means a modification of the readout from the perceptional information stored in the reservoir. This sort of tuning may happen relatively quickly within an individual’s lifetime, as in the critical learning period. Alternatively, the readout may be altered over evolutionary time by typical variation and selection processes. The puzzles concern how this happens physiologically and genetically.

Third, larger networks improve performance. Larger networks also tend to have greater redundancy with regard to storing information about the environment. Redundancy enhances robustness, provides opportunity for greater complexity, and alters evolutionary dynamics in many interesting ways [21]. This perspective raises many interesting questions about the orgin and evolution of perception.

## 4. Conclusions

Random perceptual networks may solve the puzzle of how two-step perception-response traits evolve. If a response can build on a random perceptual reservoir, then the initial evolutionary path requires adaptation only on the response side. Subsequent refinement may modify the perceptual side, changing random aspects of the initial network into more highly structured forms.

Studying the origin of traits can be difficult because we rarely observe such origins directly. Synthetic biology may provide a way to gain some insight and to test specific hypotheses. If technology advances sufficiently, it may be possible to create various types of biochemical networks that have random properties with respect to specific adaptive functions [22]. One could then use experimental evolution to analyze conditions under which cells can improve their ability to read the information in the random biochemical reservoir to achieve those specific functions.

Comparative biology could provide insight into the historical pathways and modifications of perception-response pairs. But it is not clear how easily one could find traces of evolutionary historical sequence among extant organisms. The great variety of single-cell microbial life is both promising and challenging.

## Institutional Review Board Statement

Not applicable

## Informed Consent Statement

Not applicable

## Data Availability Statement

Julia source code for the analysis and figures is available on GitHub via Zenodo at https://doi.org/10.5281/zenodo.7485008.

